# A Comprehensive Evaluation of the Genetic Architecture of Sudden Cardiac Arrest

**DOI:** 10.1101/235234

**Authors:** Foram N. Ashar, Rebecca N. Mitchell, Christine M. Albert, Christopher Newton-Cheh, Jennifer A. Brody, Martina Muller-Nurasyid, Anna Moes, Thomas Meitinger, Angel Mak, Heikki Huikuri, M. Juhani Junttila, Philippe Goyette, Sara L. Pulit, Raha Pazoki, Michael W. Tanck, Marieke T. Blom, XiaoQing Zhao, Aki S. Hauvlinna, Reza Jabbari, Charlotte Glinge, Vinicius Tragante, Stefan A. Escher, Aravinda Chakravarti, Georg Ehret, Josef Coresh, Man Li, Ronald J. Prineas, Oscar H. Franco, Pui-Yan Kwok, Thomas Lumley, Florence Dumas, Barbara McKnight, Jerome I. Rotter, Rozenn N. Lemaitre, Susan R. Heckbert, Christopher J. O’Donnell, Shih-Jen Hwang, Jean-Claude Tardif, Marja-Leena Kortelainen, Martin VanDenburgh, Andre G. Uitterlinden, Albert Hofman, Bruno H.C. Stricker, Paul I.W. de Bakker, Paul W. Franks, Jan-Hakan Jansson, Folkert W. Asselbergs, Marc K. Halushka, Joseph J. Maleszewski, Jacob Tfelt-Hansen, Thomas Engstrom, Veikko Salomaa, Renu Virmani, Frank Kolodgie, Arthur A.M. Wilde, Hanno L. Tan, Connie R. Bezzina, Mark Eijgelsheim, John D. Rioux, Xavier Jouven, Stefan Kaab, Bruce M. Psaty, David S. Siscovick, Dan Arking, Nona Sotoodehnia, for the SCD working group of the CHARGE Consortium

## Abstract

**Background:** Sudden cardiac arrest (SCA) accounts for 10% of adult mortality in Western populations. While several risk factors are observationally associated with SCA, the genetic architecture of SCA in the general population remains unknown. Furthermore, understanding which risk factors are causal may help target prevention strategies.

**Methods:** We carried out a large genome-wide association study (GWAS) for SCA (n=3,939 cases, 25,989 non-cases) to examine common variation genome-wide and in candidate arrhythmia genes. We also exploited Mendelian randomization methods using cross-trait multi-variant genetic risk score associations (GRSA) to assess causal relationships of 18 risk factors with SCA.

**Results:** No variants were associated with SCA at genome-wide significance, nor were common variants in candidate arrhythmia genes associated with SCA at nominal significance. Using cross-trait GRSA, we established genetic correlation between SCA and (1) coronary artery disease (CAD) and traditional CAD risk factors (blood pressure, lipids, and diabetes), (2) height and BMI, and (3) electrical instability traits (QT and atrial fibrillation), suggesting etiologic roles for these traits in SCA risk.

**Conclusions:** Our findings show that a comprehensive approach to the genetic architecture of SCA can shed light on the determinants of a complex life-threatening condition with multiple influencing factors in the general population. The results of this genetic analysis, both positive and negative findings, have implications for evaluating the genetic architecture of patients with a family history of SCA, and for efforts to prevent SCA in highrisk populations and the general community.

## INTRODUCTION

Sudden cardiac arrest (SCA) is a major cause of cardiac mortality, affecting over 300,000 people in the US every year^1^. Clinical and autopsy studies have demonstrated a predominant, common pathophysiology in Western populations: the most common electrophysiologic mechanism for SCA is ventricular fibrillation (VF) and the most common pathologic substrate is coronary artery disease (CAD). Despite recent increases in SCA survival rates^2^, survival remains low, and an important way to impact SCA mortality is through risk stratification and prevention. Although observational studies have identified numerous clinical and subclinical risk factors for SCA, understanding which of these associations are causal will help target prevention strategies.

Family history of SCA is a strong risk factor for SCA in the general population, suggesting that genetic variation may influence SCA risk.^3-5^ While patients with inherited arrhythmias (e.g. Long QT Syndrome) are at increased SCA risk^6-8^, the vast majority of SCA occurs outside of this high-risk population. Whether common variation in ion channel genes or other genomic regions influences SCA risk and identifies those at higher risk remains largely unknown.

Examining the genomic architecture of SCA allows us not only to examine genomic risk markers for SCA, but also to assess causal relationships of clinical and subclinical risk factors with SCA. Mendelian randomization methods exploit the fact that genetic variants are largely determined at conception and randomly distributed in populations, to determine whether an exposure may be causally associated with the outcome, and to estimate the effect size of that causal association^9-11^. Here we use a multi-SNP genetic risk score association (GRSA) model to compare genetic associations of known SCA risk factors to genetic associations with SCA as an effective way to understand the potential underlying causal pathways and processes that modulate SCA risk.

To determine whether genetic variants are associated with SCA risk, we performed a GWAS for SCA. We additionally examined whether common variation in inherited arrhythmia genes was associated with SCA risk in the general population. We then evaluated the relationships between SCA and multi-SNP GRSAs for each risk factor.

## METHODS

### Study Population and Phenotype Definition

We conducted a two-stage study. Nine studies of European-descent individuals (3,939 cases and 25,989 non-cases) comprised the GWAS ‘discovery’ stage and 12 studies with individuals of European, African and Asian descent (4,918 additional cases and 21,873 controls) comprised the ‘replication’ stage. Study descriptions, along with study-specific SCA definitions and genotyping methods, are detailed in the **Supplementary Appendix**. All studies were approved by appropriate local institutional review boards.

### GWAS

Genome-wide genotype data was imputed to the HapMap2-CEU reference panel, following study-level quality control checks (**Table S1A**). Each ‘discovery’ study performed regression analysis adjusted for age, sex, and study-specific covariates, and results were meta-analyzed using inverse variance meta-analysis implemented in METAL^12^. Metaanalysis was performed with results from 9 GWASs comprising a total of 3,939 European-ancestry cases and 25,989 controls (**Table S1A**), with additional genotyping of 26 SNPs in up to 4,918 cases and 21,879 controls of European, African, and Asian descent (**Table S1B**). For SNP rs1554218, ARIC samples were not included in the discovery data leaving 3,815 cases and 17,107 controls for the discovery stage. These ARIC samples were used only in the replication data resulting in 5,218 cases and 35,957 controls for the replication stage for analysis involving this SNP only. The top 26 SNPs were examined in a ‘replication’ population (**Table S1B**). Findings from ‘discovery’ and ‘replication’ stages were then meta-analyzed (**Table S2, Fig. S1A**). Additionally, exploratory GWASs restricted to men; women; individuals under age 65; and cases with VF/shockable rhythm, were performed (**Table S3, Fig. S1B-S1E).**

### Candidate genes

Using results from the GWAS meta-analysis, we examined variants in 54 inherited arrhythmia genes using the ‘logistic-minsnp-gene-perm’ function in FASTv1.8^13^. This best single-SNP permutation based p-value is corrected for gene size by performing up to 1 million permutations per gene. Gene boundaries were defined by RefSeq gene coordinates on build GRCh37 with +/−10 kb flanking sequence.

### Mendelian Randomization Instrument

Observational studies examine association of an exposure (e.g., body mass index, or BMI) with an outcome (e.g., SCA) but cannot assess causality. Unobserved variables affecting both exposure and outcome may confound these associations and lead to biased estimates of association. Mendelian randomization is based on the assumption that because genetic variants are determined at conception and are randomly distributed in large populations, they are unassociated with potential confounders. Therefore, under certain assumptions such as the absence of genetic pleiotropy, genetic variants used as instrumental variables can determine whether an exposure is potentially causally associated with the outcome, and estimate the size of that association (see **Supplemental Appendix**). Here we use a multi-SNP genetic risk score association (GRSA) model to compare genetic associations with SCA with those of known SCA risk factors as an effective way to understand the underlying causal pathways and processes that influence SCA risk.

### Genetic Risk Score Association (GRSA)

We estimated a separate GRSA for each of the following: (1) CAD and traditional CAD risk factors, including type 2 diabetes (T2D), fasting glucose adjusted for BMI (FGadjBMI), fasting insulin adjusted for BMI (FIadjBMI), diastolic blood pressure (DBP), systolic blood pressure (SBP), total cholesterol (TCH), and triglycerides (TG); (2) cardiac electrophysiologic factors, including atrial fibrillation (AF), heart rate (HR), QRS interval (QRS), and QT interval (QT); and (3) anthropometric traits, including BMI, waist circumference adjusted for BMI (WCadjBMI), waist to hip ratio adjusted for BMI (WHRadBMI), and height. **Table S4** details the 18 traits, and the source published GWAS used to construct the GRSA models for these traits.

To estimate GRSAs for each putative SCA risk factor, we examined genome-wide SNPs associated with the risk trait following stringent LD-pruning (**Supplementary Appendix**). The associations of these SNPs with the risk factors and the SCA outcome are used to calculate an inverse-variance weighted multi-SNP GRSA as implemented in the R-package ‘gtx’^14^. This GRSA can be interpreted as an inverse-variance weighted, meta-analyzed (over SNPs) estimate of the causal log odds ratio for SCA associated with a one SD higher value of the risk factor from a Mendelian randomization analysis.^15^ It is computationally equivalent to the slope estimate from a zero-intercept linear regression with log odds ratio for the association of an additional variant allele in SNPs with SCA (β_SCA_) as the dependent variable and the mean difference associated with one additional variant allele in SNPs on the risk factor trait (β_trait_) as the independent variable, weighted by the standard error of the β_SCA_ squared (SEsca^2^) (**Fig. 1A**) (more details in **Supplementary Appendix**). The validity of this analysis requires that SNPs included can only affect the outcome through their effects on the risk factor (i.e. no horizontal pleiotropy). If there is no pleiotropy, the SNPs contributing the GRSA estimate should all estimate the same magnitude causal association between risk factor and SCA. We use the HEIDI-outlier method from the ‘gsmr’ R package to detect and remove potentially pleiotropic SNPs.^16^ Note that we report GRSA estimates from analyses only including SNPs that meet a stringent genome-wide significant (GWS) *P*-value cut-off (*P*<5×10^-8^), GRSA_GWS_, as SNPs at this significance level likely are true positives and reliable instruments. However, the power for Mendelian randomization is dependent on the variance explained by the SNPs included in the GRSA, and for complex traits, the majority of the true signals may lie in SNPs that do not meet genome-wide significance. Therefore, we identified a somewhat arbitrary *P*-value cut-off based on visual inspection of the variance explained plots that largely maximizes variance explained while minimizing the number of SNPs (**Figure S2**). We found that all the traits fell between 0.2-0.4 *P*-value cutoff, but the results within a trait were robust to cutoffs chosen between 0.2 and 0.4. We use a GRSA constructed with this custom *P*-value cut-off (GRSA_max_) to assess only the significance of the GRSA (*P*_max_), as this model has the greatest power to assess the significance of an association. *P*_max_ is determined by permutation due to inflated test statistics (**Figure S3** and **Supplementary Appendix).** At less stringent *P*-values, false-positive SNPs may be included resulting in a bias of the estimate toward the confounded association level. Therefore, we do not use the GRSA_max_ to determine the magnitude of the GRSA association, only its direction and significance. We performed two analyses, one using GRSA_GWS_ to evaluate significance and effect size, and secondarily using the GRSA_max_ to evaluate potential associations and directions of effect at maximal power (*P*_max_). We performed multiple-testing adjustment on all resulting *P*-values (*P*_gws_ and *P*_max_) using a false discovery rate (FDR) cutoff of FDR<0.05.

**Figure 1.**
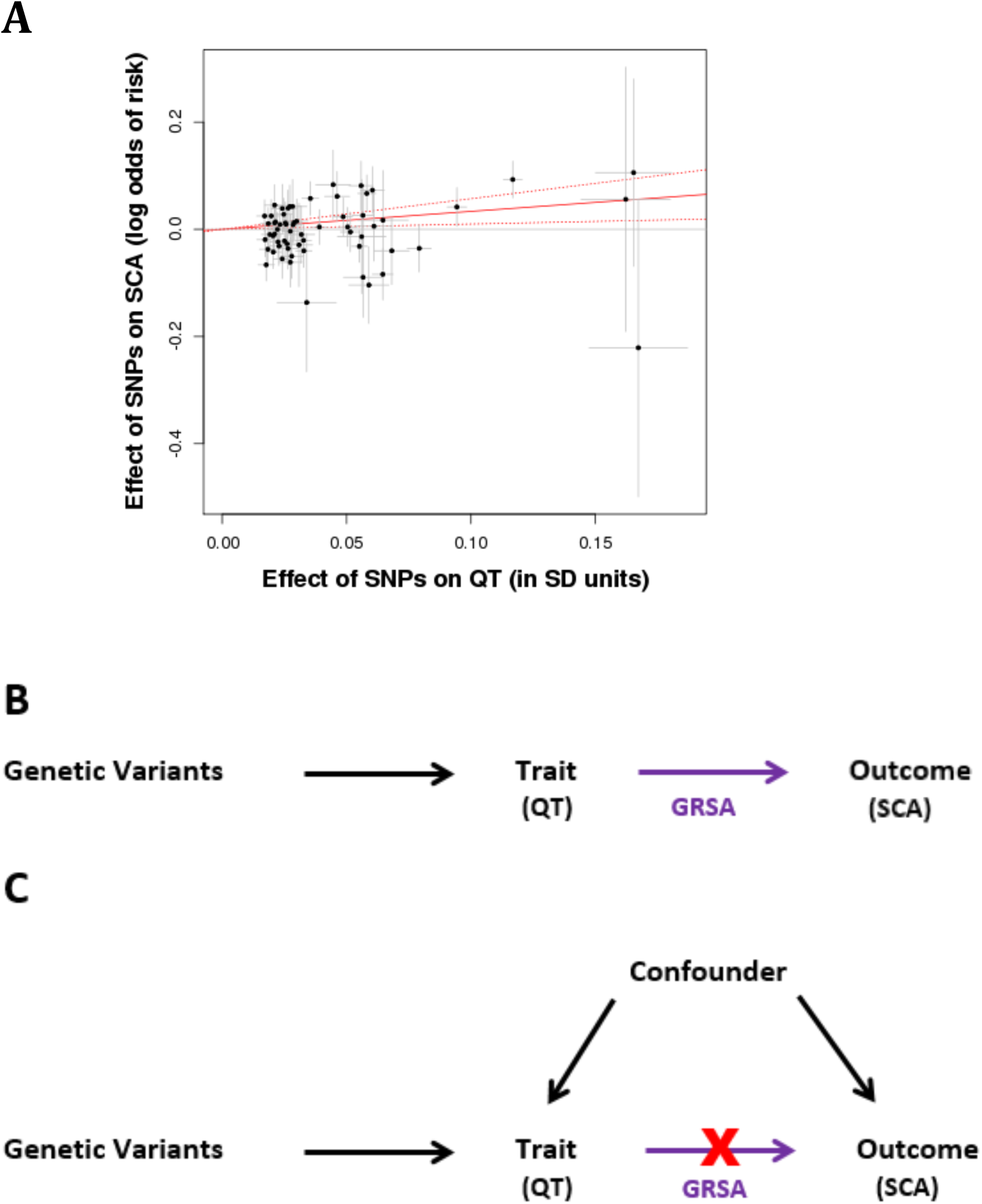
Genetic Risk Score Association (GRSA) Estimation. The plot (A) illustrates the process by which the QT-SCA GRSA is calculated using SNPs associated with QT at *P*<5×10^-8^. The points represent the effect of each SNP on QT (in units of standard deviation of QT) on the x-axis, and the log odds effect on SCA risk (corresponding 95% confidence intervals in grey) on the y-axis. The estimate of the genetic risk score association is the slope of the zero-intercept weighted regression line (solid red line). For the GRSA used in our analyses, the model contains a genome-wide LD-pruned SNP set (details in Methods). The top directed acyl graph (B) represents a scenario in which the trait of interest has a causal effect on the outcome. If the GRSA, comprised of trait-associated variants (e.g., QT), has a significant effect on the outcome (e.g., SCA), it supports a causal role for the trait on the outcome. The bottom directed acyl graph (C) presents the case where an association is observed between the trait and outcome, but the GRSA comprised of trait-associated variants is not significantly associated with the outcome, suggesting that the observational association is likely being mediated by a confounding variable and the trait does not have a causal impact on the outcome.

We similarly computed risk factor GRSAs on the outcome of CAD. We use a 1-degree of freedom Wald test to test for difference in GRSA_gws_ magnitudes between SCA and CAD.

### Sex-specific analyses

We performed sex-specific SCA GWAS analyses to construct trait GRSAs separately by sex. GRSAs were constructed from the same set of LD-pruned SNPs used for overall GRSA_gws_ analyses. *P*-values for difference in GRSA_gws_ between sexes were obtained from 1-degree of freedom Wald test.

## RESULTS

### GWAS

Meta-analysis was performed with results from 9 GWASs of 3,939 European-ancestry cases and 25,989 controls (**Table S1A, Fig. S1A**) with additional genotyping of 26 SNPs in up to 4,918 cases and 21,879 controls of European, African, and Asian descent (**Table S1B**). No SNPs were associated with SCA (*P*<5×10^-8^) (**Table S2**) in the main analysis or in subgroup analyses limited to European-descent individuals, men, women, younger participants (≤65 years), or cases with documented VF/shockable rhythm (**Tables S2** and **S3, Fig. S1B-S1E**).

### Candidate Gene and Candidate SNP Analyses

Despite sufficient power to detect relative risks of 1.15 (80% power, allele frequency 0.30, at alpha=0.05, after Bonferroni correction for multiple-testing) in a candidate gene analysis, we did not find common variants in inherited arrhythmia genes associated with SCA in the general population (**Table S5)**. Examining SNPs previously associated with SCA in smaller studies, 5/19 were nominally associated with SCA (*P*<0.05), though the current GWAS samples are not independent of those used in previous publications (**Table S6)**.

### Genetic Risk Scores Associations (GRSAs)

To explore whether clinical and subclinical risk factors are causally linked with SCA, we examined genetic risk score associations (GRSA) between SCA and: (1) CAD and traditional CAD risk factors; (2) cardiac electrophysiologic factors; and (3) anthropometric traits.

#### CAD and CAD risk factors

Prevalent CAD is an important SCA risk factor with ∼80% of male SCA survivors having underlying CAD^17^. From GRSA_GWS_ analysis we show that the difference in CAD status is causally associated with SCA (odds ratio in SCA risk per log odds difference in CAD, 1.36; 95% CI, 1.19-1.55; *P*_gws_=5.16×10^-6^) (**Fig. 2, Table S7**). While traditional CAD risk factors (blood pressure, lipids and diabetes) were not significantly associated with SCA at the more restrictive GRSA_GWS_ threshold, using GRSA_max_ to maximize power, several additional associations were detected, including type 2 diabetes (*P*_max_<0.001), LDL (*P*_max_=0.005), total cholesterol (*P*_max_<0.001), triglycerides (*P*_max_<0.001), diastolic blood pressure (*P*_max_=0.0170), and systolic blood pressure (*P*_max_=0.0230) (**Table S8)**. In the GRSA_max_ analysis, variants associated with higher diabetes risk, higher cholesterol and triglyceride levels, and higher systolic and diastolic blood pressure were all associated with higher SCA risk.

**Figure 2.**
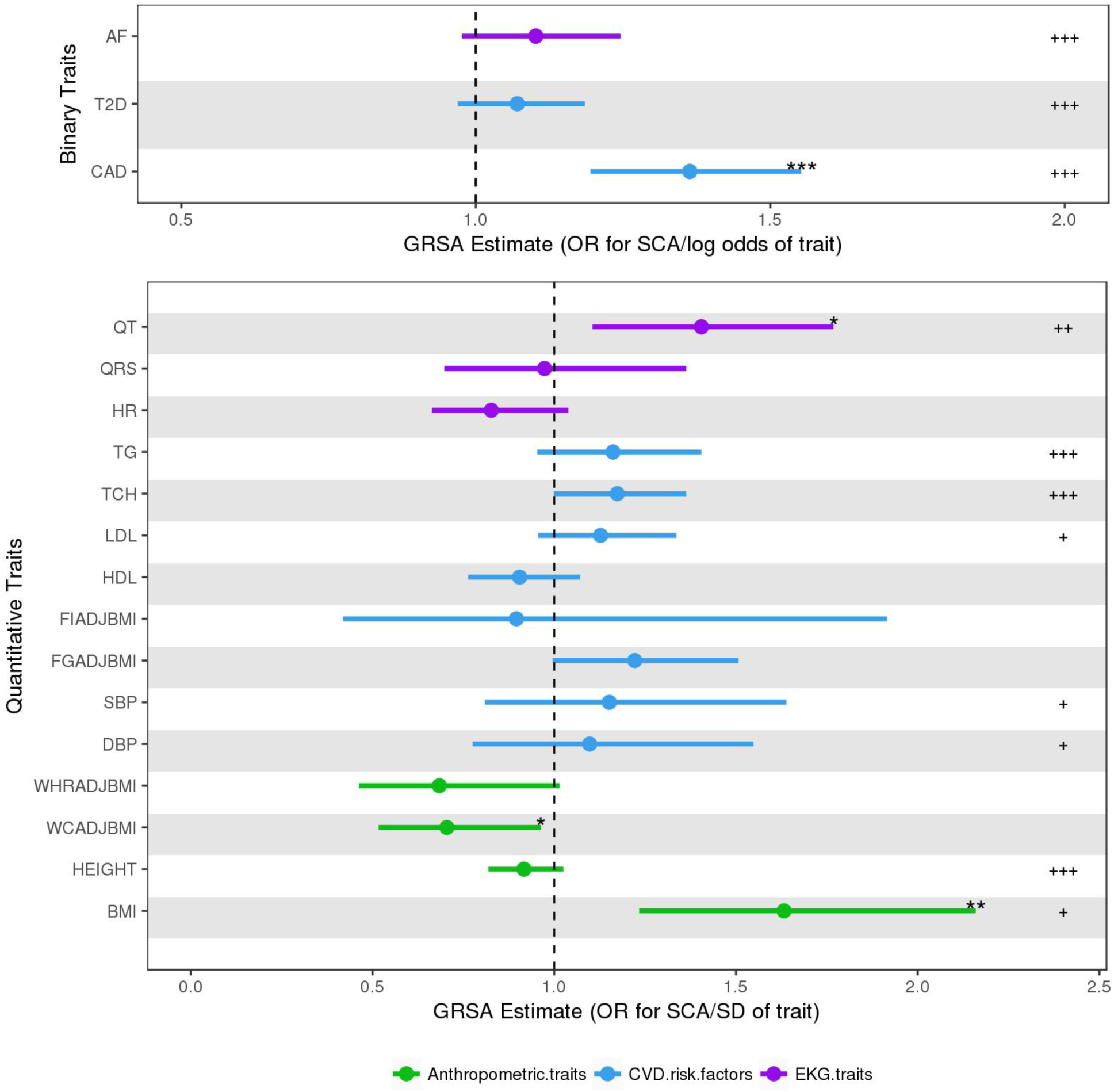
Genetic Risk Scores Association (GRSA) Estimates for SCA. These data points represent the exponentiated GRSA estimates of 18 traits on sudden cardiac arrest (SCA) and corresponding 95% confidence interval values. The GRSA estimates in the top panel for the binary traits are in log odds units. Values in bottom panel are in SD units of the quantitative traits. GRSA estimates and significance are derived from SNPs associated with each trait at *P*<5×10^-8^. The significance of the GRSA_GWS_ estimates (FDR adjusted *P*_GWS_) are represented as “*” for *P*<0.05, “**” for *P*<0.01, and “***” for *P*<0.001. The significance of the secondary analysis using GRSA_max_ estimates (FDR adjusted permuted *P*_max_) are represented as “+” for *P*<0.05, “++” for *P*<0.01 and “+++” for *P*<0.001. For details on values of GRSA estimates and *P*-values, see **Table S7-S8.** CAD = coronary artery disease; T2D = type 2 diabetes; AF = atrial fibrillation; BMI = body mass index; WCadjBMI = waist circumference adjusted for BMI; WHRadBMI = waist to hip ratio adjusted for BMI; DBP = diastolic blood pressure; SBP = systolic blood pressure; FGadjBMI = fasting glucose adjusted for BMI; FIadjBMI = fasting insulin adjusted for BMI; HR = heart rate; QRS = QRS interval; QT = QT interval; HDL = high-density lipoproteins; LDL = low-density lipoproteins; TCH = total cholesterol; TG = triglycerides.

#### Cardiac electrophysiologic factors

To explore the influence of cardiac electrophysiology on SCA, we examined genetics of electrophysiologic traits associated with SCA: (1) atrial fibrillation, (2) QT interval (ventricular repolarization), (3) QRS interval (ventricular conduction), and (4) heart rate. In the GRSA_GWS_ analysis, we show that longer QT interval, a risk factor for SCA in the general population, is significantly associated with SCA (odds ratio in SCA risk per SD increase in QT, 1.40; 95% CI, 1.10-1.77; Pgws=0.005) **(Fig. 2, Table S7)**.^18^ Using GRSA_max_, in addition to QT, we also identified a significant association of AF with SCA (*P*_max_<0.001 for both QT and AF) (**Table S8**). Variants associated with longer QT interval and higher AF risk were associated with higher SCA risk. By contrast, no significant association was seen with QRS or heart rate, even at the more permissive and statistically powerful GRSA_max_.

#### Anthropometric Measures

The BMI GRSA_GWS_ was significantly associated with SCA (odds ratio for SCA risk per SD higher BMI, 1.63; 95% CI, 1.23-2.15; *P*_GWS_=6.02×10^-4^) (**Fig. 2, Table S7).** Using GRSA_max_, we found a significant negative association between height and SCA (*P*_max_<0.001) **(Table S8)**. Variants associated with greater height are associated with lower CAD risk^19^, and we correspondingly observed a negative GRSA between SCA and height. No significant association was seen with GRSAs composed of variants associated with measures of central/abdominal adiposity, such as waist-to-hip ratio or waist circumference.

### Contrasting SCA and CAD GRSAs

Given the strong association of CAD with SCA, we compared the magnitudes of risk factor GRSA_gws_ on the outcomes of SCA (**Fig. 2**) and CAD (**Fig. S4**) to identify traits where risk factors may be more strongly causally associated with SCA than CAD. While the GRSA_gws_ for traditional CAD risk factors (blood pressure and lipid traits) are larger for CAD risk than SCA risk, we find that GRSA_gws_ for electrophysiologic traits of QT interval (0.34 for SCA vs. 0.096 for CAD, *P* for difference = 0.06) and AF (0.097 for SCA vs. −0.029 for CAD, *P* for difference=0.017), there was a suggestion of a larger association with SCA than CAD risk (**Fig. 3, Table S7**).

**Figure 3.**
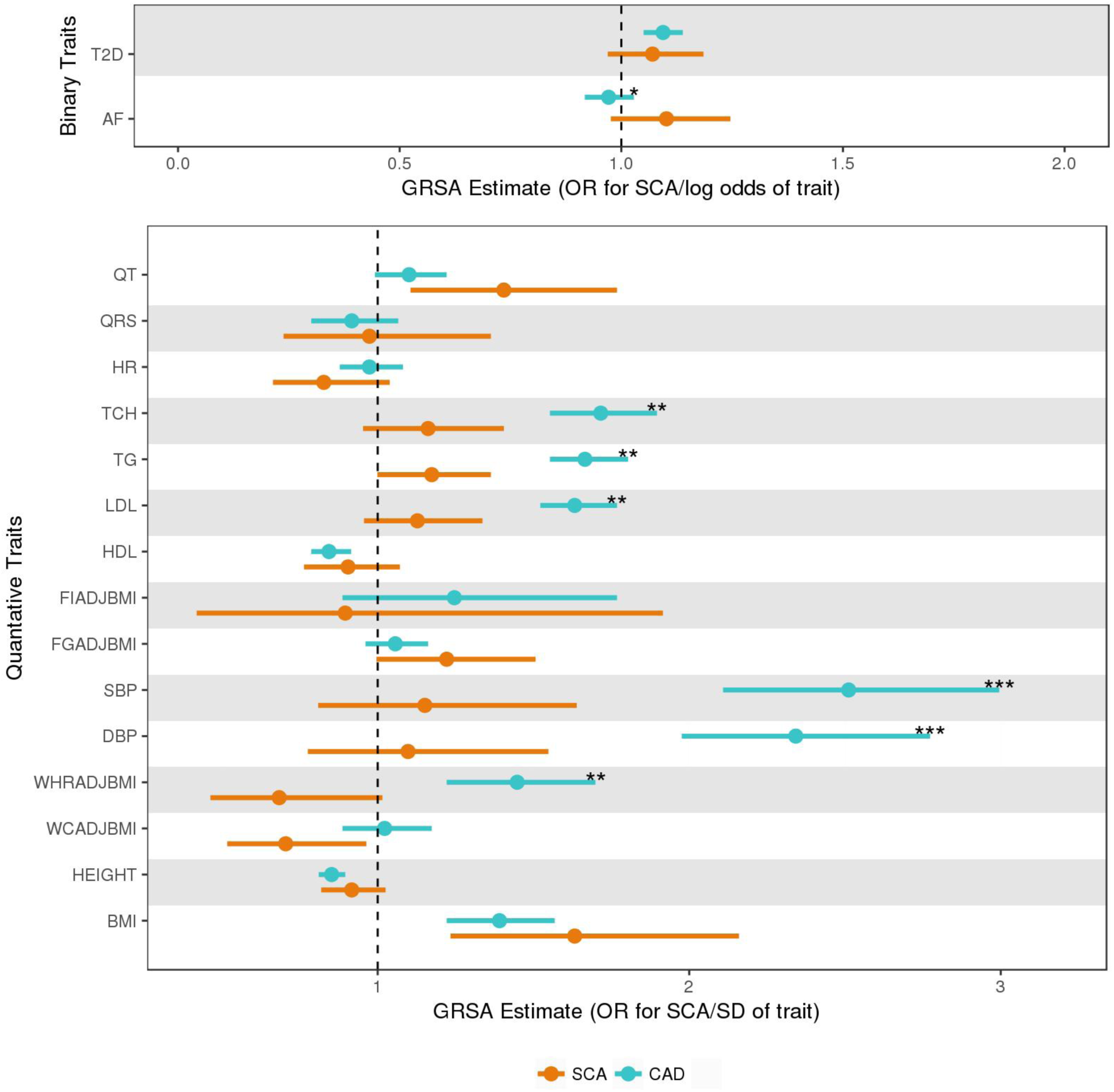
Comparison of GRSA for SCA and CAD. These data represent exponentiated GRSAs of all 17 traits. GRSA estimates for SCA and CAD, are plotted in orange and teal respectively. Bars around the estimates represent the 95% confidence interval. The GRSA estimates in the top panel for the binary traits are in log odds units. Values in bottom panel are in SD units of the quantitative traits. The level of significance for 1 degree of freedom Wald test of difference in GRSA_gws_ estimates between SCA and CAD is represented “*” for *P*<0.05, “**” for *P*<0.01, and “***” for *P*<0.001. T2D = type 2 diabetes; AF = atrial fibrillation; BMI = body mass index; WCadjBMI = waist circumference adjusted for BMI; WHRadBMI = waist to hip ratio adjusted for BMI; DBP = diastolic blood pressure; SBP = systolic blood pressure; FGadjBMI = fasting glucose adjusted for BMI; FIadjBMI = fasting insulin adjusted for BMI; HR = heart rate; QRS = QRS interval; QT = QT interval; HDL = high-density lipoproteins; LDL = low-density lipoproteins; TCH = total cholesterol; TG = triglycerides.

### Sex differences

Sex differences in SCA incidence, underlying SCA pathophysiology, and prevalence of certain risk factors have been well documented^19^, yet little is known about whether the effect of risk factors on SCA differs by sex. Among GRSAsgws where a main effect association was identified, we found a nominally significant difference in association between women and men for diabetes (0.240 for women vs. 0.0205 for men, *P* for difference = 0.05) and HDL (-0.417 for women vs. 0.0256 for men, *P* for difference = 0.04) (**Table S9**).

## DISCUSSION

Our SCA GWAS demonstrates that while SCA is a complex disease with multiple risk factors, a comprehensive genetic approach can shed light on causal versus correlational associations. Using Mendelian randomization, we establish that differences in CAD, BMI, and QT interval are causally associated with SCA. Secondary analyses further implicate type 2 diabetes, additional traditional CAD risk factors such as lipids and blood pressure, as well as height and atrial fibrillation.

Despite adequate power to identify relatively modest associations, our study did not find evidence that common variation in Mendelian arrhythmia genes is associated with SCA risk in the general population. Since underlying electrical instability is an important cause of SCA, prior smaller studies have examined inherited arrhythmia genes or variants associated with electrophysiological traits to identify genetic variants that influence SCA risk ^21-23^. While rare private mutations in ion-channel and other electrophysiology-related genes increase arrhythmia risk in high-risk families and may also increase SCA risk in the general population^24^, our study suggests that common variants in these genes are not significant contributors to SCA in the general population. This may be due to differing underlying genetics between inherited arrhythmias versus SCA in the general population. By contrast, we do find that GRSA estimates of phenotypes associated with electrical instability (AF and QT) are causally associated with SCA risk, more so than they are causally associated with CAD. This confirms our understanding of the pathophysiology of SCA—SCA is not simply fatal CAD, but rather, electrical instability also plays a prominent role in influencing SCA risk.

Intriguingly, not all electrophysiologic phenotypes observationally linked to SCA are causally associated with SCA in our analyses. QRS interval and heart rate, two traits observationally associated with SCA^25^,^26^, failed to show significant evidence of a shared genetic basis with SCA. This lack of association may be due to inadequate power to identify more modest correlations. Alternatively, it may be that the associations from observational studies are confounded by other factors, and not causative **(Fig. 1B-C)**. For instance, underlying CAD can lead to both longer QRS interval and increased SCA risk; thus, while observational studies show an association between SCA and both traits (CAD and QRS interval), the association between SCA and QRS interval may not be causal. Similarly, the observational association of higher heart rate with SCA risk may be confounded by higher adrenergic state due to underlying heart disease and not itself be causal. Thus, the GRSA approach to examining observational risk factors assists in differentiating causative factors from confounded associations.

CAD is the most common underlying pathologic substrate for SCA. It is reassuring, therefore, that we find significant estimated causal associations with SCA risk using GRSA models constructed from CAD and traditional CAD risk factors, including blood pressure, diabetes and cholesterol traits.

Anthropometric measures appear to be causally associated with SCA. Shorter stature is associated with increased SCA risk in observational studies; our findings support the conclusion that this observational association is causal. Observational data on BMI and SCA risk have been conflicting, perhaps due to confounding from smoking status and frailty. Previously^27^, we have shown that increased BMI is associated with increased SCA risk in non-smokers, but not smokers. In this study, we find that differences in BMI, but not central/abdominal obesity, were causally associated with SCA risk. This finding is especially interesting in the context of recent data that imply different biological process underlying BMI and central obesity.^28^,^29^

Finally, of the traits associated with SCA, we found that GRSAs for diabetes and HDL were nominally significantly different between men and women. While diabetes is a SCA risk factor among both sexes, previous observational studies have consistently suggested a stronger, albeit not statistically different, association among women than men^30^,^31^. These findings may reflect different underlying SCA pathophysiology between men and women. While these differences may be due to chance as they do not remain significant after multiple test correction, it is also likely that our study is underpowered to detect these differences.

Several limitations deserve consideration. First, without detailed autopsy information, rhythm monitoring, and information on circumstances surrounding the cardiac arrest, the underlying etiology and mechanism of death may be heterogeneous and genetic associations are likely to be diluted. Nonetheless, clinical and autopsy studies have demonstrated a predominant, common pathophysiology of SCA in Western populations: VF in the setting of CAD. Hence, it is reassuring that our genetic studies suggest an important role for both CAD and electrical instability in SCA. Second, despite ours being the largest exploration of SCA genomics to date, the discovery sample size of only ∼4,000 cases limited our ability to find genetic associations with low frequency variants or variants of modest effect. Hence, while our data do not support screening individuals with a family history of SCA for common variation in inherited arrhythmia genes, much larger samples sizes are needed to address whether rare variation of modest effect in these genes influence SCA risk. Third, the validity of the GRSA method as a Mendelian randomization instrument rests on the assumption that the variant causes differences in the outcome only by its effects on the risk factor of interest, and not directly or by influencing other risk factors. Although we did not explicitly exclude SNPs associated with multiple risk factors (genetic pleiotropy), we did utilize a goodness-of-fit approach to exclude putative “pleiotropic” effects from all GRSAs. Furthermore, while genetic pleiotropy can bias our conclusions, important influence is less likely when using multiple SNPs aggregated in a genetic risk score.

In conclusion, while we were not able to identify any common genetic variants significantly associated with SCA risk through the GWAS, as well as any common variation in specific inherited arrhythmia genes associated with SCA risk, we have provided evidence for causal associations between some, but not all, observational risk factors for SCA. We show that differences in CAD status, BMI, and QT interval are causally associated with SCA risk. While SCA is a complex disease with multiple influencing factors, a comprehensive genetic approach can untangle risk factor relationships, enhancing our understanding of SCA pathophysiology. Ultimately, genetic studies will enhance efforts to prevent SCA in high-risk populations and the general community.

### Disclosures

Bruce M. Psaty serves on the DSMB of a clinical trial funded by Zoll LifeCor and on the Steering Committee of the Yale Open Data Access Project funded by Johnson & Johnson.

### Funding Support

#### AGNES

This study was supported by research grants from The Netherlands Heart Foundation (grants 2001D019, 2003T302 and 2007B202), the Leducq Foundation (grant 05-CVD), the Center for Translational Molecular Medicine (CTMM COHFAR) and the Netherlands Heart Foundation (CVON project PREDICT). C.R.B. is an Established Investigator of The Netherlands Heart Foundation (grant 2005T024) and is supported by the Netherlands Organization for Scientific Research (NWO, grant ZonMW Vici Project number: 016.150.610).

#### ARREST

This study has received funding from the European Union’s Horizon 2020 research and innovation programme under acronym ESCAPE-NET, registered under grant agreement No 733381. H.L.T. is supported by The Netherlands Organization for Scientific Research (NWO, grant ZonMW Vici 918.86.616) and the Medicines Evaluation Board Netherlands (CBG/MEB).

#### ARIC

The Atherosclerosis Risk in Communities Study is carried out as a collaborative study supported by National Heart, Lung, and Blood Institute contracts (HHSN268201100005C, HHSN268201100006C, HHSN268201100007C, HHSN268201100008C, HHSN268201100009C, HHSN268201100010C, HHSN268201100011C, and HHSN268201100012C), R01HL087641, R01HL59367 and R01HL086694; National Human Genome Research Institute contract U01HG004402; and National Institutes of Health contract HHSN268200625226C. Infrastructure was partly supported by Grant Number UL1RR025005, a component of the National Institutes of Health and NIH Roadmap for Medical Research. FNA, AM, and DEA were supported by R01HL11267 and R01HL116747.

#### CABS

The Cardiac Arrest Blood Study is supported by the following grants from the National Heart, Lung, and Blood Institute (NHLBI): HL111089, HL088456, HL088576, HL091244, HL116747, and HL092111. Additional funding was provided by the Laughlin Family and the John Locke and Medic One Foundations.

#### CARTAGENE

This study is supported by the French National Institute of Medical and Scientific Research (INSERM), Direction Générale de la Santé (French ministry of Health) and a French German grant BMBF-ANR for the genetic analyses.

#### CHS

This study was supported by NHLBI contracts HHSN268201200036C, HHSN268200800007C, N01HC55222, N01HC85079, N01HC85080, N01HC85081, N01HC85082, N01HC85083, N01HC85086; and NHLBI grants U01HL080295, R01HL087652, R01HL105756, R01HL103612, R01HL120393, and R01HL130114 with additional contribution from the National Institute of Neurological Disorders and Stroke (NINDS). Additional support was provided through AG023629 from the National Institute on Aging (NIA). A full list of CHS investigators and institutions can be found at http://www.chs-nhlbi.org/pi.htm. The provision of genotyping data was supported in part by the National Center for Advancing Translational Sciences, CTSI grant UL1TR001881, and the National Institute of Diabetes and Digestive and Kidney Disease Diabetes Research Center (DRC) grant DK063491 to the Southern California Diabetes Endocrinology Research Center. The content is solely the responsibility of the authors and does not necessarily represent the official views of the National Institutes of Health. NS was supported by HL111089, HL116747, and the Laughlin Family Endowment.

#### FHS

This study is supported by National Heart, Lung, and Blood Institute grant HL-54776 and contracts 53-K06-5-10 and 58-1950-9-001 from the USDA Research Service.The Framingham Heart Study SHARe Project was partially supported by the NHLBI Framingham Heart Study (Contract No. N01-HC-25195) and its contract with Affymetrix, Inc for genotyping services (Contract No. N02-HL-6-4278). DNA isolation and biochemistry were partly supported by NHLBI HL-54776. A portion of this research utilized the Linux Cluster for Genetic Analysis (LinGA-II) funded by the Robert Dawson Evans Endowment of the Department of Medicine at Boston University School of Medicine and Boston Medical Center.

#### FINGESTURE

This cohort is supported by the Juselius Foundation (Helsinki, Finland), the Council of Health of the Academy of Finland (Helsinki, Finland), and the Montreal Heart Institute Foundation.

#### GEVAMI

This study was funded by grant supports from the University of Copenhagen (Copenhagen, Denmark), The Danish National Research Foundation (Copenhagen, Denmark), The John and Birthe Meyer Foundation (Copenhagen, Denmark), The Research Foundation of the Heart Center Rigshospitalet (Copenhagen, Denmark), and The Bikuben Scholar-Danmark-Amerika Fonden & Fulbright Commission (Copenhagen, Denmark).

#### KORAF3

This study was initiated and financed by the Helmholtz Zentrum München - German Research Center for Environmental Health, which is funded by the German Federal Ministry of Education and Research (BMBF) and by the State of Bavaria. KORA research was additionally supported within the Munich Center of Health Sciences (MC-Health), Ludwig-Maximilians-Universität, as part of LMUinnovativ.

#### HARVARD

This study was supported by grants: HL-092111, HL-068070, HL-26490, HL-34595, HL-34594, HL-35464, HL-46959, HL-080467 from the National Heart, Lung, and Blood Institute and CA-34944, CA 40360, CA55075, CA-87969, CA 97193 from the National Cancer Institute.

#### ROTTERDAM

This study is funded by Erasmus Medical Center and Erasmus University, Rotterdam, Netherlands Organization for the Health Research and Development (ZonMw), the Research Institute for Diseases in the Elderly (RIDE), the Ministry of Education, Culture and Science, the Ministry for Health, Welfare and Sports, the European Commission (DG XII), and the Municipality of Rotterdam. The generation and management of GWAS genotype data for the Rotterdam Study is supported by the Netherlands Organisation of Scientific Research NWO Investments (nr. 175.010.2005.011, 911-03-012). This study is funded by the Research Institute for Diseases in the Elderly (014-93-015; RIDE2), the Netherlands Genomics Initiative (NGI)/Netherlands Organisation for Scientific Research (NWO) project nr. 050-060-810.

## ACKNOWLEDGEMENT

The authors thank all the staff and participants from the studies contributing to this manuscript for their important contributions.

## AGNES

We thank L. Beekman for technical support and N. Bruinsma for assistance in collection of AGNES subject data, and Marie Cécile Perier and the Emergency medical mobile units which participated in the CARTAGENE study and the genetic INSERM Unit of François Cambien in Paris, particularly Carole Proust.

